# Age-Related Differences in Alpha Power for Distractor Inhibition During Visual Working Memory

**DOI:** 10.1101/2023.06.28.546988

**Authors:** Sabrina Sghirripa, Lynton Graetz, John G. Semmler, Rebecca Sutton, Ellen E.R. Williams, Mitchell R. Goldsworthy

**Author notes:** **Corresponding author:** Sabrina Sghirripa.

## Abstract

As visual working memory (WM) is limited in capacity, it is important to direct neural resources towards task-relevant information and away from task-irrelevant information. Neural oscillations in the alpha frequency band (8-12 Hz) have been suggested to play a role in the inhibition of distracting information during WM retention in younger adults, but it is unclear if alpha power modulation also supports distractor inhibition in older adults. Here, we recorded electroencephalography (EEG) while 24 younger (aged 18-35) and 24 older (aged 60-86) adults completed a modified delay match-to-sample task in which distractors of varying strength appeared during the retention period. We found: (1) strong distractors impaired WM performance compared with weak and no distractors in both age groups, but there were no age-differences in WM performance; (2) while younger adults demonstrated significant increases in alpha power prior to the onset of the distractor in all conditions, decreases in alpha power were seen in all distractor conditions in older adults; (3) there was no difference in alpha power between the strong and no distractor conditions; and (4) alpha power in anticipation of the distractor was only associated with task performance in younger adults. Our results suggest that younger adults, but not older adults, modulate alpha power in anticipation of distractors during the visual WM retention period.

## 1 Introduction

Visual working memory (WM) is severely limited in capacity (Cowan, 2001), highlighting the importance of encoding and retaining task-relevant, and ignoring task-irrelevant or distracting information. An inability to inhibit distracting information is one suggested basis for age-related reductions in WM performance (Hasher & Zacks, 1988), but the neural mechanisms underlying age-related deficits in ignoring distractions during WM are unclear.

Neural oscillations in the alpha (8-12 Hz) frequency range have been implicated in distractor inhibition during WM performance in younger adults (Bonnefond & Jensen, 2012; Sauseng et al., 2009; Sghirripa et al., 2020). While it was initially thought that increases in alpha power reflected ‘cortical idling’ (Pfurtscheller et al., 1996), the modulation of alpha power has now been proposed to dynamically gate sensory input to task-relevant brain regions (Jensen & Mazaheri, 2010). Evidence linking alpha power to distraction inhibition during visual WM was derived from tasks where participants attend to a task-relevant and inhibit a task irrelevant hemifield. These paradigms are associated with decreases in alpha power in the task-relevant, and increases in alpha power in the task-irrelevant hemifield, implicating alpha power in both the facilitation of visual WM performance and inhibition of task-irrelevant information (Sauseng et al., 2009).

During the retention period of verbal WM tasks where participants can anticipate the onset of a distractor, anticipation of strong distractors has been associated with greater alpha power prior to the onset relative to alpha power in anticipation of a weak distractor (Bonnefond & Jensen, 2012). Using a verbal WM paradigm, we have previously reported an increase in alpha power that did not differ based on distractor type (Sghirripa et al., 2020), while others using visual WM tasks have reported lower alpha power in the presence of strong distractors present for the entire retention period (Schroeder et al., 2018).

Alpha oscillations reduce in power and frequency with age (Babiloni et al., 2006), and given that age-related deficits in distractor inhibition may account for age-related deficits in WM performance (Hasher & Zacks, 1988), a link may exist between age-related changes in alpha oscillatory activity and deficits in distractor inhibition during WM. In a study employing a lateralised task to investigate age-differences in alpha power for the suppression of irrelevant information during WM, it was observed that while suppression of visual processing was associated with modulation of alpha power in younger adults, older adults did not modulate alpha power, despite performance indicating that suppression of distractors had occurred (Vaden et al., 2012).

However, older adults demonstrate greater performance declines when distractors match the same category of stimulus as the memory set, suggesting that the strength of the distractor is important in age-related declines in distractor inhibition (Clapp et al., 2009; Clapp & Gazzaley, 2012; Yoon et al., 2006). Despite studies in younger adults reporting differences in alpha power modulation with distractor strength (Bonnefond & Jensen, 2012), no study has investigated whether older adults also modulate alpha power in anticipation of distractors of varying strength, or whether alpha power modulation is absent in older adults, as described by Vaden et al. (2012).

Here, we employed a modified delay match-to-sample task with strong, weak and no distractor conditions to determine whether young and older adults modulate alpha power in anticipation of distractors of varying strength. We hypothesised that: 1) WM performance would be more impaired by strong and weak distractors in older, compared with younger adults, 2) that younger adults would show stronger alpha power before the onset of strong, compared with weak distractors, while older adults would show no differences in alpha power between conditions, and 3) that stronger alpha power in the lead up to the distractor would correlate with better WM performance.

## 2 Method

### 2.1 Participants

24 healthy younger (mean age: 23.36 years, SD: 5.83 years, range:18-35, 22 female) and 24 healthy older adults (mean age: 69.96 years, SD: 6.94 years, range: 60-86, 17 female) participated in the study. The samples in each group were not significantly different for years of education (*t*_47_ = 0.50, *p* = 0.62). All older adults were without cognitive impairment as assessed by Addenbrooke’s Cognitive Examination III (ACE-III) (score > 82) (Mioshi et al., 2006).

Exclusion criteria involved a history of neurological/psychiatric disease, use of central nervous system altering medications, history of alcohol/substance abuse and uncorrected hearing/visual impairment.

All participants gave informed written consent before the commencement of the study, and the experiment was approved by the University of Adelaide Human Research Ethics Committee.

### 2.2 Working Memory Task

Participants first completed a WM load adjustment task consisting of 20 trials at each of load-2, load-3, load-4, load-5 and load-6 with no distractors present. This task trained the participant on the task and allowed us to adjust the WM load for each participant individually before partaking in the distractor visual WM task with EEG. The load for each participant was chosen based on the load where ∼80% accuracy was achieved.

The distractor visual WM task used stimuli presented by PsychoPy (Peirce, 2007). Each trial began with the participant fixating on a cross in the centre of the screen for 2 s. A memory set consisting of six squares then appeared for 0.5 s to maintain equal sensory input for each WM load. The position of squares did not change within trials but varied between trials. The number of coloured squares depended on the WM load chosen in the load adjustment task. Each square subtended 0.65° x 0.65° of visual angle. Following a consolidation period of 0.25 s, a pattern mask consisting of coloured static appeared for 0.15 s in the same location as each item presented in the memory set. We chose to include a mask and consolidation period to disrupt residual sensory trace left by the memory set from influencing oscillatory power during the retention period (Woodman & Vogel, 2005). Following the mask, a 4 s retention period began. In the distractor blocks, 2 s into the retention period a strong (4 coloured squares not in the memory set) or weak distractor was shown (4 light grey squares) for 0.5 s, followed by the 1.5 s remainder of the retention period. In the no distractor condition, only the fixation cross was present for 4 s. A single probe coloured square then appeared in the location of a randomly selected square from the memory set. Participants were instructed to respond with the right arrow key if the probe square was the same colour in the memory set and respond with the left arrow key if the square was a different colour. In each block, 50% of the trials required a right arrow key response (same colour), and 50% required a left arrow key response (different colour). The probe remained on the screen until the subject responded.

In the task, there were 5 blocks with strong distractors, 5 blocks with weak distractors and 5 blocks with no distractor. Blocks consisted of 20 trials (total = 100 trials per distractor condition), with the blocked design allowing participants to anticipate the strength of the distractor within blocks.

Distractors were never part of the memory set or probe, and participants were explicitly asked to ignore the distractor. Short breaks were allowed between blocks.

### 2.3 EEG Data Acquisition

EEG data were recorded from a 64-channel cap containing Ag/AgCl scalp electrodes arranged in a 10–10 layout (Waveguard, ANT Neuro, Enschede, The Netherlands) using a Polybench TMSi EEG system (Twente Medical Systems International B.V, Oldenzaal, The Netherlands). Conductive gel was inserted into each electrode using a blunt-needle syringe to reduce impedance to < 5 kΩ. The ground electrode was located at AFz. Signals were amplified 20x, online filtered (DC-553 Hz) and sampled at 2048 Hz. Due to the lack of data from the mastoids, data were referenced to the average of all electrodes. EEG was recorded during each block of the distractor visual WM task.

### 2.4 EEG Data Pre-Processing

EEG data were pre-processed using EEGLAB (Delorme & Makeig, 2004), TMS-EEG Signal Analyser (TESA v1.1.1) (Rogasch et al., 2017) and custom scripts using MATLAB (R2020b, The Mathworks, USA). Each block of EEG data was merged, and incorrect trials were flagged for removal at a later stage. Unused channels were removed, data were downsampled to 256 Hz and then band-pass (0.1-100Hz) and band-stop (48-52 Hz) filtered using the EEGLAB ‘eegfiltnew’ function. Data were epoched -2 to 6 s relative to the onset of the memory set. Channels and trials were then visually inspected and removed if contaminated with residual artifacts (e.g. muscle activity or non-stereotypical artifacts). An average of 1 channel was removed from the younger adult group, and 2 were removed from the older adult group (range young: 0-7, range old: 0-6). Independent component analysis (ICA) was then completed using the FastICA algorithm (Hyvärinen & Oja, 2000), with the ‘symmetric approach’ and ‘tahn’ contrast functions selected. Components corresponding to eye-blinks and persistent muscle activity were detected using the TESA (Rogasch et al., 2017) ‘compselect’ function and were manually inspected before removal from the data. Missing channels were then interpolated, and data were re-referenced to the common average. Epochs were then split into distractor types, and correct and incorrect trials were separated.

For younger adults, an average of 76 trials for the no distractor condition, 76 for the weak distractor condition and 72 for the strong distractor condition were accepted for further analysis. For older adults, an average of 73 trials for the no distractor condition, 72 for the weak distractor condition and 67 for the strong distractor condition were accepted for further analysis.

### 2.5 Spectral Analysis

Spectral analysis was performed using FieldTrip toolbox (Oostenveld et al., 2011). Data were converted to the time-frequency domain using a multi-taper transformation based on multiplication in the frequency domain (cfg.method = ‘mtmconvol’). A time window of 3 cycles was used for each frequency (0.5 Hz steps between 3 and 45 Hz) and time point (50 ms steps). Spectral power was calculated for individual trials before being averaged over trials for each distractor condition in each age group. Data were baseline corrected -1 to -0.25 s relative to the onset of the memory set.

### 2.6 Statistical Analysis of Behavioural Data

Statistical analyses were performed using R version 4.1.0. Linear mixed effects models with age group and distractor condition as fixed effects, participant as random effect, and accuracy and RT as outcome variables, were used to analyse behavioural data. Bonferroni corrected post-hoc tests were performed in the case of significant main effects or interactions. Correlations between the difference in alpha power between strong and weak distractor conditions and difference in accuracy and RT between conditions were performed using Pearson’s correlation. Data were checked for normality using Shapiro-Wilk tests, and the residuals for the linear mixed effects models were checked via QQ plots and histograms. In all tests, a p-value of < 0.05 was considered statistically significant. Data are presented as mean ± SEM in figures.

### 2.7 Statistical Analysis of EEG Data

Statistical analyses of EEG data were performed using FieldTrip toolbox (Oostenveld et al., 2011). Cluster-based permutation tests were used to assess differences in alpha power between age-groups and distractor types. Cluster-based permutation tests control for the type-1 error rate when comparing across multiple channels, times, and frequencies (Maris & Oostenveld, 2007). Clusters were defined as two or more neighbouring electrodes for which the difference in spectral power between age groups (independent samples t-test) or distractor types (dependent samples t-test), exceeded *p* < 0.05. A permutation distribution was used to test clusters for significance, which was generated by combining alpha power values from both age groups and distractor types into a single set, randomly partitioning into two subsets, and taking the largest cluster-level statistic from this partition (Monte Carlo method; 2000 random permutations). If the cluster-level statistic observed from the original data was larger in absolute value than 95% of random partitions, the cluster was deemed significant (*p* < 0.05, two-tailed test).

To test for interactions between age group and distractor type, we employed a 2×2 factorial design. A difference power spectrum was calculated consisting of the power spectrum differences between distractor types for each combination of no, weak, and strong distractor (e.g., for differences between strong and weak distractors: young difference = Young/Strong - Young/Weak, Old difference = Old/Strong - Old/Weak). Cluster-based permutation tests performed as described above to compare the difference power spectrums between age groups.

## 3 Results

### 3.1 Behavioural Data

#### 3.1.1 WM Load Adjustment

In the younger adult group, 9 participants completed the task at load-4 and 15 completed the task at load-5. In the older adult group, 8 participants completed the task at load-3, 13 completed the task at load-4 and 3 completed the task at load-5. The load that was selected for each participant was significantly different between groups (*t*_46_ = 4.96, *p* < 0.001), with younger adults (M = 4.63, SD = 0.50) being able to perform near 80% accuracy at higher loads than older adults (M = 3.79, SD = 0.66).

#### 3.1.2. RT

For RT there were significant main effects of age group (*F*_1,46_ = 35.84, *p* < 0.001) and distractor type (*F*_2,84_ = 15.62, *p* < 0.001), but no age group by distractor type interaction (*F*_2,84_ = 0.08, *p* = 0.92). Bonferroni corrected post-hoc tests revealed that older adults were slower to respond in all conditions than younger adults (*p* <0.001). Response times were slower in the strong distractor condition compared with the no (*p* < 0.001) and weak (*p* = 0.005) distractor conditions, but there was no difference in response times between the weak and no distractor conditions (*p* = 0.23) (Figure 1B).

**Figure 1.**
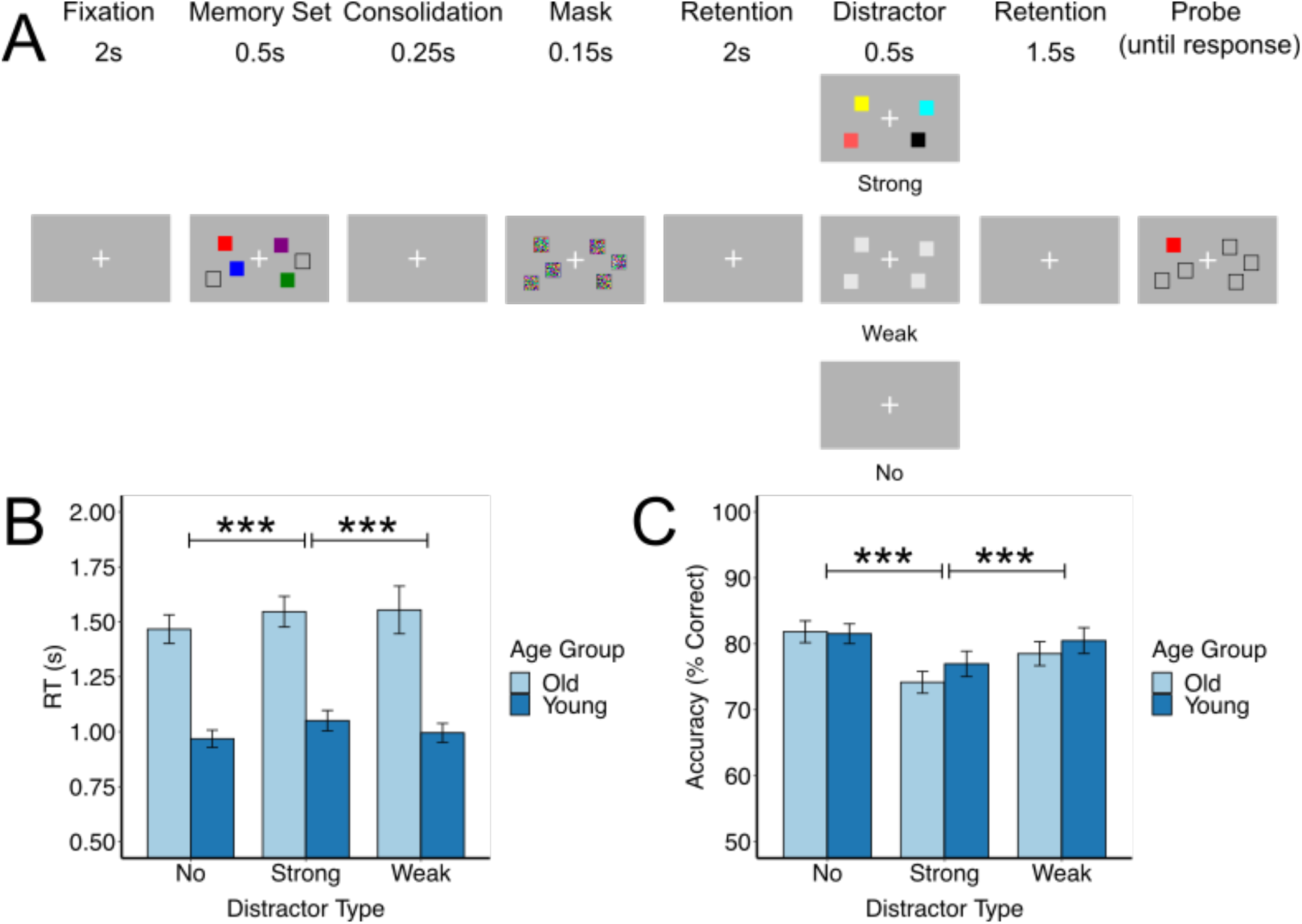
A) Modified delay match-to-sample task. Each trial contained a memory set with a number of squares chosen via performance on the load adjustment task with no distractors present, followed by a pattern mask after a short delay. The retention period was 4 s long, in which a strong (4 coloured squares) or weak (4 grey squares) distractor was shown and persisted for 0.5 s. In the no distractor blocks, only a fixation cross was present for the 4 s retention period. A single probe square was then presented, and participants responded to whether the coloured square was the same colour seen in the memory set. B) Reaction time (RT) for correct responses to the probe and C) accuracy (% correct) in response to the probe in each distractor condition *** *p* <0.001.

#### 3.1.3 Accuracy

For accuracy there was a significant main effect of distractor type (*F*_2,84_ = 23.67, *p* < 0.001), but no main effect of age (*F*_1,46_ = 1.34, *p* = 0.25), nor an age group by distractor type interaction (*F*_2,84_ = 1.72, *p* = 0.19). Bonferroni corrected post-hoc tests revealed that accuracy was poorer in the strong distractor condition compared with the no (*p* < 0.001) and weak (*p* < 0.001) distractor conditions, but there was no difference in accuracy between the weak and no distractor conditions (*p* = 0.38) (Figure 1C).

### 3.2 Time Frequency Analysis

Our a-priori hypothesis was that increases in alpha power would be present prior to the onset of distractors. When we examined the time-frequency representation of raw power averaged across both age groups and all distractor conditions from all participants, and the average of all electrodes, the largest alpha power was observed in the 0.75 s prior to the onset of the distractor in the 9-12 Hz frequency range. In the cluster-based permutation tests, we averaged across the 9-12 Hz frequency range and the 0.75 s prior to distractor onset and tested across all electrodes.

#### 3.2.1 Age Differences in Anticipatory Alpha Power

We sought to determine whether age differences in alpha oscillatory power existed in the pre-distractor time interval for each distractor type. Upon visual inspection of the time-frequency representations of power, we observed that alpha power increased from baseline in all distractor conditions in younger adults, whereas alpha power decreased from baseline across all conditions in older adults (Figure 2A). In younger adults, the increase in alpha power from baseline was significant for the no distractor (*p* = 0.01) and strong distractor (*p* = 0.04) conditions, but not for the weak distractor condition (no significant clusters). The decrease in alpha power from baseline was significant in older adults for the no distractor (*p* = 0.01), weak distractor (*p* = 0.002) and strong distractor conditions (*p* = 0.04).

**Figure 2.**
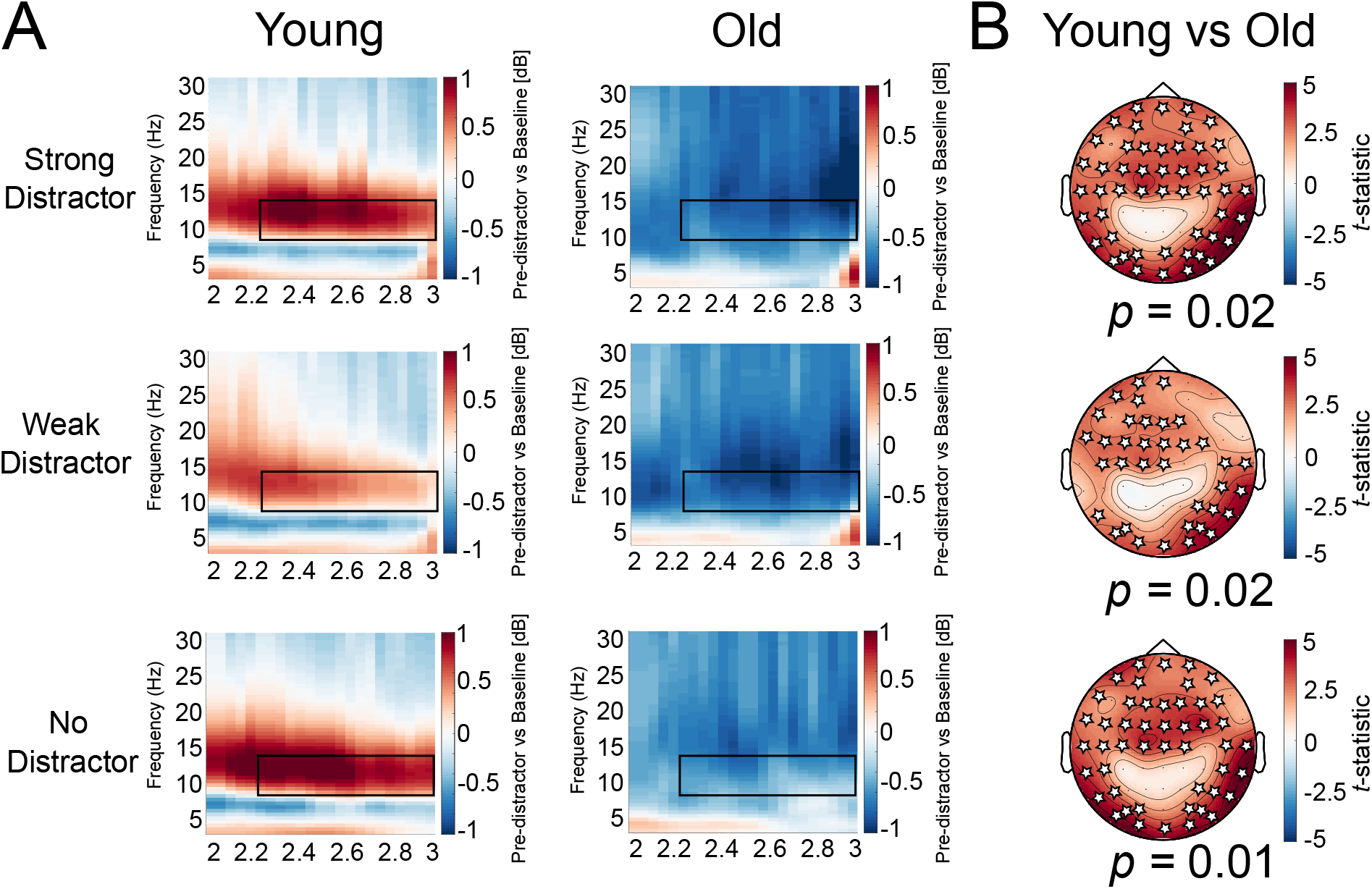
A) Baseline corrected time-frequency representations of power (average of all electrodes) in the 1 s prior to the onset of the distractor, and B) t-statistics from cluster-based permutation tests demonstrating the difference between age-groups in each distractor condition. Black boxes in time-frequency plots indicate times and frequencies of interest for cluster-based permutation tests, and white stars indicate electrodes in significant clusters.

Cluster based permutation tests revealed that during anticipation of a distractor, younger adults demonstrated larger increases in alpha power in the strong (*p* = 0.02) and weak (*p* = 0.02) distractor conditions compared with older adults. However, we also found age differences in alpha power in the no distractor condition (*p* = 0.01). In all comparisons, the differences in alpha power were prominent across frontal, parietal, and occipital electrodes (Figure 2B).

#### 3.2.2 Differences Between Distractor Types

We then sought out to determine whether alpha power prior to the distractor differed between distractor conditions in both younger and older adults.

In younger adults, cluster-based permutation tests revealed a significant increase in alpha power in the strong, relative to weak distractor condition in the 9-12 Hz range in the 0.75 s preceding the distractor (*p* = 0.03). These differences were seen in the left frontal and frontocentral, and right parietal, parieto-occipital and occipital electrodes. In contrast, cluster-based permutation tests revealed no significant difference between strong and weak distractor types in older adults (all *p* > 0.07) (Figure 3A). However, we could not find evidence for a distractor type by age group interaction (no significant clusters) (Figure 3B).

**Figure 3.**
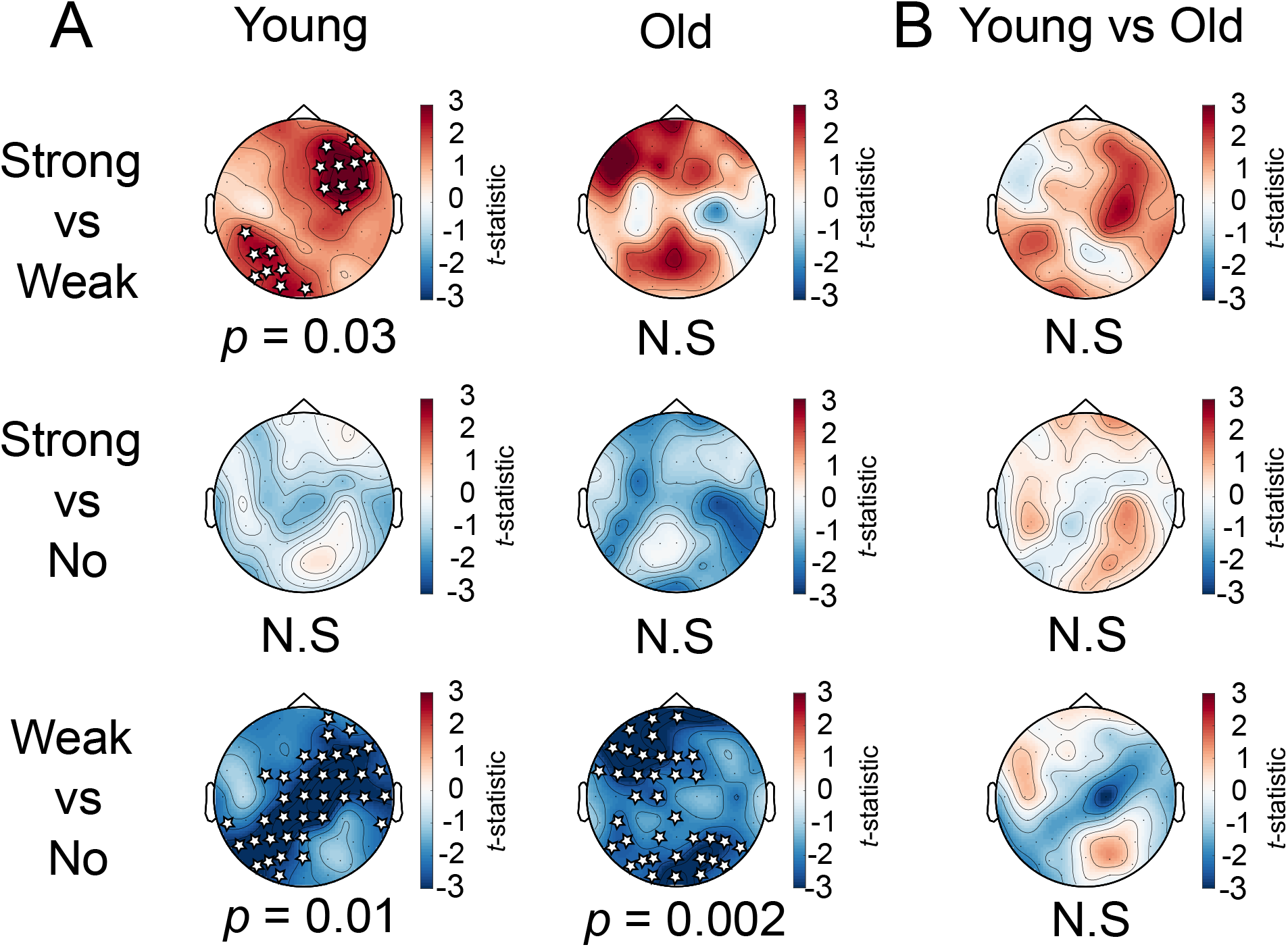
A) *t*-statistics for cluster-based permutation tests comparing the differences in alpha power prior to the distractor for each distractor type. B) t-statistics for cluster-based permutation tests for an interaction effect between age-groups and the difference between distractor types. White stars indicate electrodes in significant clusters.

**Figure 4.**
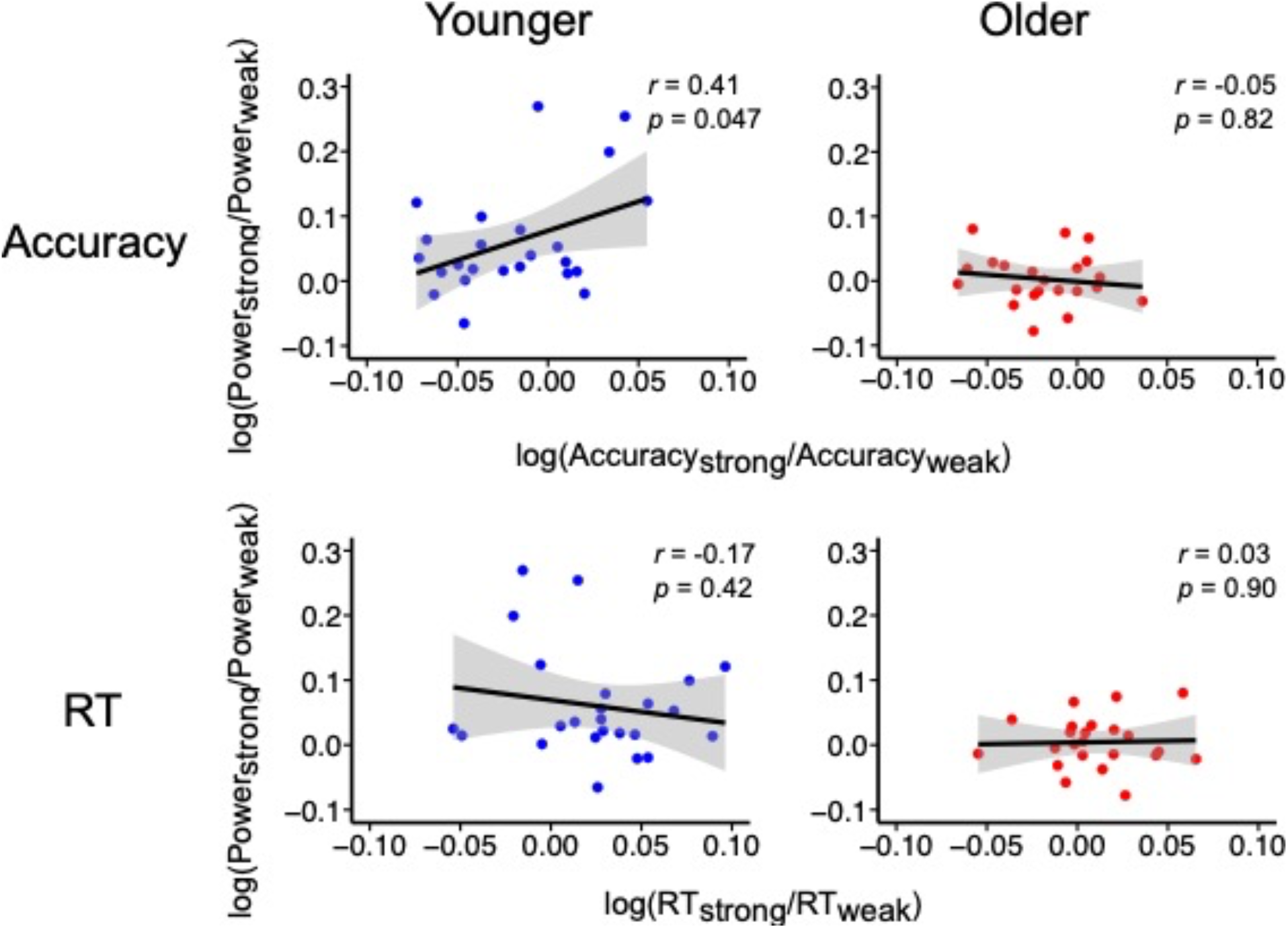
Correlations between the difference in alpha power and difference in accuracy (top) and RT (bottom) for younger (left) and older (right) adults.

We then investigated whether there were differences in alpha power between each distractor condition and the no distractor condition. In both younger and older adults, cluster-based permutation tests revealed no significant differences in alpha power between the no distractor condition and the strong distractor condition (no significant clusters for young, *p* = 0.08 for old). In both younger and older adults, alpha power was lower in the weak distractor, compared with the no distractor condition (younger adults: *p* = 0.01, older adults: *p* = 0.002).

#### 3.3.3 Alpha Power and Task Performance

First, we compared correct with incorrect trials from each participant for each distractor condition to determine whether there were differences in alpha power in trials with a correct compared with an incorrect response. Cluster-based permutation tests revealed no significant differences between correct and incorrect trials in any distractor conditions, across each age group (all *p* > 0.07).

To determine whether alpha power prior to the distractor influenced task performance, we conducted a correlation between the difference in alpha power in the 0.75 s before the onset of the distractor (data derived from electrodes in the significant cluster for younger adults, and from all electrodes for older adults) and the difference in accuracy and RT between strong and weak distractor trials. We found a moderate correlation between accuracy and alpha power in younger adults (*r* = 0.41, *p* = 0.047), indicating that participants with stronger difference in alpha power between distractor types demonstrated greater accuracy. However, we did not find a correlation between the difference in alpha power and RT (*r* = -0.17, *p* = 0.42) in younger adults, nor a correlation between the difference in alpha power and difference in accuracy (*r* = 0.05, *p* = 0.82) or difference in RT (*r* = -0.03, p = 0.9) between distractor types in older adults.

## 4 Discussion

In this study, we investigated age-related differences in alpha oscillatory power in anticipation of distractors of varying strength during the visual WM retention period. Contrary to our hypothesis, we found that when WM load was individualised to each participant, there was no difference in accuracy between age groups in any distractor condition. Despite the lack of behavioural difference, we found that while younger adults demonstrated significant increases in alpha power from baseline prior to the onset of the distractor in the strong and no distractor conditions, decreases in alpha power from baseline were seen in all distractor conditions in older adults. When comparing alpha power between distractor types, younger adults demonstrated greater alpha power before the onset of strong, relative to weak distractors, while older adults demonstrated no difference in alpha power between distractor types. However, there was no difference in alpha power between the strong distractor and no distractor conditions, and surprisingly, alpha power was greater in the no distractor, compared with the weak distractor condition. Finally, alpha power in anticipation of strong distractors was moderately correlated with WM performance in younger adults.

### 4.1 Strong distractors impaired WM performance, but distractor inhibition does not differ between age groups

In this study, we could not find evidence for age-related deficits in distractor inhibition during WM. Regardless of age group, we found that strong distractors impaired WM performance compared with both the weak and no distractor conditions. Although older adults were slower to respond to the probe in each distractor condition and completed the task at a lower WM load on average, we did not find age differences in the cost of distraction.

An absence of age differences is broadly inconsistent with extensive literature detailing the inhibitory deficit hypothesis, which suggests that older adults experience WM deficits due to an inability to inhibit task-irrelevant information (Hasher & Zacks, 1988). Our results are also inconsistent with previous work that has demonstrated that age-related decline in distractor inhibition during visual WM is greater for distractors occurring in the retention period as opposed to those presented in the encoding period (McNab et al., 2015).

However, our results align with the findings of Vaden et al. (2012), who also demonstrated no age-differences in WM performance in the presence of distractors. One explanation for our lack of age difference is that like Vaden et al. (2012), the predictability of the distractor both in temporal onset and strength may have assisted the older adults with ignoring the distractor. Previous work has shown that both distraction (task-irrelevant stimuli) and interruptions (stimuli that must be attended to as a secondary task) negatively impact WM performance, but interrupters disproportionately affect WM performance in older adults (Clapp & Gazzaley, 2012). The fact that our task only required participants to ignore the distractor rather than respond to it in some way may have been less taxing on executive control systems that are affected by age-related decline (Clapp et al., 2009). Likewise, many studies investigating suppression of task-irrelevant information have used complex stimuli such as faces and scenes (Clapp & Gazzaley, 2012; Gazzaley & D’esposito, 2007), which may be harder to inhibit than simple stimuli like coloured squares.

Finally, we recruited participants for this study via convenience sampling. Convenience sampling of older adults generally results in the recruitment of participants who perform better cognitively than older adults in the wider population (Brodaty et al., 2014), which may explain the lack of age difference seen here.

### 4.2 Younger adults, but not older adults show increases in alpha power in anticipation of distractors

In the younger adults we found that alpha power in anticipation of distractors was higher in the strong, relative to the weak distractor condition, with some evidence to suggest that higher alpha power before strong distractors is associated with better WM performance. Our results align with those of Bonnefond and Jensen (2012), who found stronger increases in alpha power in occipito-temporal areas in anticipation of strong, compared with weak distractors. However, older adults demonstrated decreases in alpha power in anticipation of distractors, consistent with the findings of Vaden et al. (2012), who also reported that despite successfully ignoring irrelevant information, older adults did not modulate alpha to suppress distractors.

Alpha oscillations have long been thought to be involved in top-down suppression of task irrelevant information during WM, in tasks both with and without distractors (Bonnefond & Jensen, 2012; Jensen et al., 2002; Jensen & Mazaheri, 2010; Klimesch, 2007). Deficits in top-down suppression have been implicated in age-related declines in WM performance (Gazzaley et al., 2005), and given the suggested role of alpha power in inhibition, it is possible that an inability to modulate alpha power plays a role in impaired suppression of irrelevant information in older adults. Although we found that older adults demonstrated a decrease in alpha power before the distractor as opposed to an increase, suggesting deficits in top-down suppression, we did not find evidence for age-related deficits in distractor inhibition, nor a link between alpha power modulation and behavioural performance in older adults. If the lack of alpha modulation seen in older adults is evidence of altered top-down suppression mechanisms, then it is possible that older adults are using an alternative neural strategy to compensate for their inability to modulate alpha power. At rest, alpha power decreases with advancing age (Babiloni et al., 2006; Lindsley, 1939), which may suggest that alpha power modulation is not a viable neural strategy for older adults to use when ignoring distractions during WM due to their existing low alpha power (Vaden et al., 2012).

Alternatively, the decrease in alpha power prior to the distractor may suggest that older adults are encoding the distractors rather than ignoring them, as decreases in alpha power have been shown to represent increased visual cortex excitability for impending stimuli (Romei et al., 2008, 2010). Older adults may be more susceptible to encoding, rather than inhibiting the distractor, as the ‘deletion’ facet of the inhibitory deficit hypothesis posits that age related declines in WM performance may be due to older adults allowing task-irrelevant information to enter the WM store, and then being unable to delete the distractions (Hasher & Zacks, 1988). However, the lack of a behavioural difference between younger and older adults suggests that even if older adults are encoding the distractors, their WM performance is not being negatively affected by the distractor, regardless of whether the content of the distractor matches the memoranda.

### 4.3 No differences in alpha power between strong and no distractors, despite a difference between strong and weak distractors in younger adults

Visual WM retention is often associated with a decrease in alpha power during the retention period, which is thought to reflect maintenance of visual information. For example, it has been shown that alpha power decreases with visual WM load and the degree of alpha suppression correlates with individual WM capacity (Fukuda et al., 2015). Given the visual nature of the task, we expected to see decreases in alpha power during the no distractor condition relative to baseline and compared with the strong and weak distractor conditions. Contrary to our hypothesis, we observed no significant difference in alpha power between the strong distractor and no distractor conditions in both young and older adults.

The results of our no distractor condition are at odds with the visual WM literature, and align with the pattern of alpha activity reported in verbal WM tasks, where increases in alpha power are commonly seen during retention (Jensen et al., 2002; Proskovec et al., 2019; Tuladhar et al., 2007; Wang et al., 2016). The increase in alpha power during the verbal WM retention period has been interpreted to reflect inhibition of the visual stream to prevent task-irrelevant information from entering the WM store, even in the absence of visual distractors, which could explain the increase in alpha power in the no distractor condition seen here. While we used a block design to clearly segregate strong and weak distractor trials from no distractor trials, presenting all three conditions in the same session could have led to participants using a similar neural strategy in all conditions, leading to a general brain state change even when distractors are not present (van Diepen & Mazaheri, 2017). Therefore, the no distractor condition should be performed in a separate experimental session to confirm whether the demands of distractor conditions induced this change in alpha power, or whether the demands of the visual WM retention period itself, led to increases in alpha power.

Additionally, many visual WM tasks using change detection paradigms that report alpha suppression during the retention period involve tasks with retention periods of approximately 1 s (Adam et al., 2017; Fukuda et al., 2015; Sghirripa et al., 2022). We have previously shown that the alpha suppression seen during retention in tasks with a ∼1 s retention period is due to residual alpha suppression from encoding (Sghirripa et al., 2022), which may not be present in longer retention periods. Although Fukuda et al. (2015) performed an experiment with a 4 s retention period and reported alpha suppression for the entirety of retention, participants performed the task at load-1 and load-3, which may not represent a degree of difficulty where participants need to employ neural strategies to prevent task-irrelevant information from entering the WM store. Therefore, more research is required to determine the pattern of alpha power modulation during visual WM, and the functional significance of the effect.

It remains unclear why the weak distractor condition resulted in lower alpha power than the no distractor condition, particularly in the younger adult group given the moderate correlation between the difference in alpha power between distractor types and task performance. A potential explanation for this is that if content of the visual distractor did not compete, or weakly competed with the active WM store, some participants may have encoded the weak distractor, resulting in decreases in alpha power due to visual expectation prior to distractor onset (Romei et al., 2008, 2010). If this is the case, encoding the weak distractor likely did not have deleterious effects on task performance, given the lack of behavioural difference between the weak and no distractor conditions in both age groups. Conversely, if the visual distractor did compete with the contents of memory, then alpha power increased to the same extent during retention as if no visual stimulus occurred, possibly reflecting anticipatory suppression of encoding to protect the contents of WM from the distractor, or from temporal decay of the WM store. Regardless, further work is required to understand the role of alpha power in distractor inhibition during WM.

### 4.4 Conclusion

Here, we found that younger adults demonstrate increases in alpha power in anticipation of distractors during the visual WM retention period and that this increase supports WM performance in the presence of strong distractors. In contrast, older adults demonstrate decreases in alpha power in anticipation of distractors. Despite the differences in alpha power modulation between age groups, we did not find evidence of age-related deficits in distractor inhibition, suggesting that older adults employ different neural strategies to inhibit distractors during visual WM. Further work should now investigate the neural mechanisms underlying distractor inhibition during WM in older adults.

